# Myosin XV is a negative regulator of signaling filopodia during long-range lateral inhibition

**DOI:** 10.1101/2023.07.07.547992

**Authors:** Rhiannon Clements, Tyler Smith, Luke Cowart, Jennifer Zhumi, Alan Sherrod, Aidan Cahill, Ginger L Hunter

## Abstract

The self-organization of cells during development is essential for the formation of healthy tissues, and requires the coordination of cell activities at local scales. Cytonemes, or signaling filopodia, are dynamic actin-based cellular protrusions that allow cells to engage in contact mediated signaling at a distance. While signaling filopodia have been shown to support several signaling paradigms during development, less is understood about how these protrusions are regulated. We investigated the role of the plus-end directed, unconventional MyTH4-FERM myosins in regulating signaling filopodia during sensory bristle patterning on the dorsal thorax of the fruit fly Drosophila melanogaster. We found that Myosin XV is required for regulating signaling filopodia dynamics and, as a consequence, lateral inhibition more broadly throughout the patterning epithelium. We found that Myosin XV is required for limiting the length and number of signaling filopodia generated by bristle precursor cells. Cells with additional and longer signaling filopodia due to loss of Myosin XV are not signaling competent, due to altered levels of Delta ligand and Notch receptor along their lengths. We conclude that Myosin XV acts to negatively regulate signaling filopodia, as well as promote the ability of signaling filopodia to engage in long-range Notch signaling. Since Myosin XV is present across several vertebrate and invertebrate systems, this may have significance for other long-range signaling mechanisms.

## Introduction

The formation of robust and reproducible patterns is essential during development. Initially unordered precursor cells can be organized to form simple spot arrays, stripes, or more complex 3-dimensional structures like intestinal villi or limbs. The incorrect spatial or temporal organization of cells during development can lead to catastrophic defects in organ function or early embryonic lethality. Generally, the patterning of cells requires input from morphogenetic cues that can be distributed in both a passive and active manner throughout the tissue. Passive processes include the short range diffusion of signaling molecules (Stapornwongkul et al., 2020). Active processes include the distribution of signaling molecules via cytonemes, or signaling filopodia (Kornberg and Roy, 2014; Zhang and Scholpp, 2019). Cytonemes are long (*>* 2μm), thin (≈200nm), actin-based projections that support the delivery of signaling molecules (Kornberg, 2017; Ramírez-Weber and Kornberg, 1999). Cytonemes are involved in numerous patterning events during the development of the fruit fly Drosophila melanogaster including: distributing Dpp/TGF*β* (Roy et al., 2014) and Hh (Bischoff et al., 2013; Chen et al., 2017; González-Méndez et al., 2020) associated with wing disc development; distribution of growth factors during mechanosensory bristle development (Peng et al., 2012); supporting long-range Notch signaling (Huang and Kornberg, 2015; Hunter et al., 2019); and Hh-mediated stem cell niche maintenance (Rojas-Ríos et al., 2012). In vertebrates, cytonemes are known to play a role in the distribution of Shh, Wnt, and Notch (Eom et al., 2015; Hall et al., 2021; Hamada et al., 2014; Mattes et al., 2018; Sanders et al., 2013; Stanganello et al., 2015) in developmental processes ranging from pigment stripe formation, neural plate formation, and limb bud morphogenesis. While active processes have been the focus of intense research across several signaling paradigms, how changes in cell morphology affect the ability of cells to send or receive signal are only beginning to be understood. There is evidence that feedback from signaling mechanisms supported by cytonemes can stimulate the formation of longer or additional cytonemes (de Joussineau et al., 2003; Snyder et al., 2015). Disruption of cytoskeletal components that regulate cell shape can disrupt signaling output in cytoneme-dependent processes (Cohen et al., 2010; Hunter et al., 2019). However, the strategies used to disrupt the cytoskeleton are usually not specific to cytonemes, and broadly affect sub-cellular processes that are dependent on actin and microtubules. Very few tools to specifically perturb cytonemes exist (Zhang et al., 2021). Further investigation is needed to gain insight into how the molecular mechanisms that regulate cell shape changes coordinate with cell-cell signaling processes.

The sensory bristle pattern on the dorsal thorax of Drosophila melanogaster is a model system for the study of pattern formation mediated by signaling filopodia (Heitzler and Simpson, 1991; Collier et al., 1996; Cohen et al., 2010; Corson et al., 2017; Hunter et al., 2019). Over several hours during pupal development, a regular and well-spaced array of sensory bristle precursor cells is formed from a sheet of bipotential epithelial cells. The selection of bristle precursor cells occurs through canonical Notch-mediated lateral inhibition (Bray, 2016; Troost et al., 2015). When transmembrane Delta ligand and Notch receptor proteins interact in trans-, the Notch receptor is activated through two proteolytic cleavages that lead to the release and translocation of the intracellular domain (NICD) to the nucleus. Once in the nucleus, the NICD plays a context dependent role on transcription with its co-factors, including Suppressor of Hairless (CBF1 or Lag-1) and Mastermind (Kopan and Ilagan, 2009). During bristle patterning, cells with low levels of Notch activation allow the expression of genes associated with a pro-neural fate, and commit to the bristle lineage. Cells with high levels of Notch activation repress the pro-neural genes and commit to an epithelial cell fate. In order to achieve the correct density of bristle precursor cells, a long-range Notch-mediated lateral inhibition signal is required (+1-2 cell diameters)(Cohen et al., 2010; Collier et al., 1996; Corson et al., 2017; Hadjivasiliou and Hunter, 2022). Long, actin-rich protrusions are formed on the basal surface of thoracic epithelial cells during patterning stages (de Joussineau et al., 2003; Cohen et al., 2010; Hunter et al., 2019; Renaud and Simpson, 2001; Georgiou and Baum, 2010). These protrusions are dynamic and allow cells to contact neighbors 2-3 cell diameters away. Several regulators of the actin cytoskeleton are required to form the basal signaling filopodia, including SCAR, Rac, cdc42, and non-muscle myosin II (Cohen et al., 2010; Georgiou and Baum, 2010; Hunter et al., 2019). Long-range contact between two or more basal signaling filopodia carrying Notch receptor or Delta ligand should lead to the activation of cell surface Notch, however, direct evidence for this mechanism has yet to be obtained.

A key factor in the generation of actin-based cell protrusions such as cytonemes are the actin binding proteins that organize the cytoskeleton. Among the actin-binding myosin motors that play a role in regulating cellular protrusions are the unconventional class of MyTH4-FERM domain containing myosins, including Myosin VIIa, VIIb, X, and XV (Sellers, 2000; Weck et al., 2017). Mutations in the human genes that encode Myosin VIIa and XV are associated with congenital forms of deafness (Hasson et al., 1995; Wang et al., 1998; Weil et al., 1995). Mammalian Myosin VIIa and Myosin XV play roles in maintaining the function of stereocilia: VIIA maintains the integrity of the tip complex to help organize the characteristic ‘staircase’ organization of inner ear hair cells (Boeda, 2002; Self et al., 1998), where as XV trafficks actin regulators to stereocilia tips that help control the structure’s length (Belyantseva et al., 2005; Manor et al., 2011). Mammalian Myosin VIIb is involved in the maintenance and formation of microvilli (Weck et al., 2016), while Myosin X is closely associated with the formation and function of filopodia (Berg and Cheney, 2002; Heimsath et al., 2017). Notably, Myosin X has also been shown to play a role in the trafficking of signaling molecules in cytonemes, as well as in the formation cytonemes (Hall et al., 2021; Snyder et al., 2015). The Drosophila melanogaster genome encodes three unconventional Myosins: VIIa, VIIb, and XV. In the fly, loss of Myosin VIIa causes deafness (Todi et al., 2005) and disrupts the formation of other actin-based projections during development (Sallee et al., 2021). It is currently unknown what effect the loss of Myosin VIIB has on cell morphology and development (encoded by the gene myosin 28B1, FBgn0040299). Loss of Drosophila Myosin XV leads to defects in cell sheet migration during embryogenesis (Liu et al., 2007), as well as defects in the elongation of sensory bristles, which require parallel bundles of actin filaments (Rich et al., 2021). Interestingly, the Drosophila genome lacks a gene that encodes Myosin X. Despite a lack of Myosin X, which supports cytoneme signaling in vertebrates, there are many morphogenetic events that rely on the activity of cytonemes and filopodia in Drosophila (Bischoff et al., 2013; Cohen et al., 2010; Huang et al., 2019; Huang and Kornberg, 2015; Hunter et al., 2019; Millard and Martin, 2008; Ramírez-Weber and Kornberg, 1999). This suggests that, in the fly, there are alternative pathways to supporting the formation of, and trafficking within, these cellular protrusions.

In this study, we investigate the mechanisms that regulate the morphology and dynamics of signaling filopodia and how these behaviors contribute to the progression of Notch-mediated spot patterning. We show that the formation of basal signaling filopodia in the notum requires the activity of Myosin XV. We further investigate the requirement for Myosin XV in the sub-cellular distribution of Notch and Delta in bristle precursor cells. To understand better the mechanisms by which Myosin XV promotes signaling filopodia length, we investigate which of the motor domains are required for full-length filopodia formation and Delta localization. Together, our data sheds light on the mechanisms that regulate the activity and formation of signaling filopodia as well as a new role for the unconventional myosin motor Myosin XV.

## Materials and methods

### Fly husbandry

Drosophila stocks were maintained on standard Drosophila food (JazzMix, Fisher Scientific), at 18°C with 24 hour light cycle. Crosses are maintained at room temperature (23-25°C). White pre-pupae were screened against balancers and then aged at 18°C for 24 hours in humidified chambers. Pupae aged to 12-14 hours after pupariation were then dissected for live or fixed imaging protocols.

### Immunofluorescence

Dissected nota of 12-14 hAP pupae were fixed for 20 minutes in 4% paraformaldehyde/1X PBS solution. Tissues were then blocked in 1:1 blocking buffer (5% w/v BSA, 3% FBS in 1X PBS) in 1X PBST at room temperature for 1 hour. Tissues were then incubated in primary antibody with 5% blocking buffer in 1X PBST for 2 hours at room temperature or 4°C overnight. Tissues were washed twice in 1X PBST for 10 minutes each at room temperature, followed by incubation with secondary antibody in 1X PBST containing fluorescently-labeled phalloidin and DAPI, for 2 hours at room temperature or 4°C overnight. Tissues were washed twice in 1X PBST for 10 minutes each, then equilibrated in 50% glycerol overnight. Nota were then mounted on coverslips and sealed with nail varnish for storage at 4°C until imaged. The following primary antibodies were used in this study: chicken anti-GFP (EDMillipore 1:1000), mouse anti-NotchECD (C458.2H, DSHB, 1:250), mouse anti-Delta (C594.9B, DSHB, 1:250). The following secondary antibodies were used in this study: AlexaFluor 488 donkey anti-chicken (Jackson Immunological, 1:2000), AlexaFluor 647 anti-mouse. The following stains were also used: ActiStain Phalloidin (Phdh1, Cytoskeleton Inc, 1:500), DAPI (Thermoscientific, 1:1000).

### Imaging

Samples were imaged on a either a Lieca DMi8 SPE confocal microscope using LASX software, or Nikon C2+ or EclipseTi confocal microscopes using NIS Elements. The pupal cases of live 12-14 hAP pupae were removed, exposing the head and thorax. A coverslip coated with a thin film of Halocarbon 27 oil (Sigma) was placed on top of spacers such that only the dorsal thorax contacted the coverslip (as previously published (Loubéry and González-Gaitán, 2014)). Live pupae were imaged using a x40 (0.8 NA) air objective (SPE) or or x60 (1.4 NA) oil objective (EclipseTi). Fixed tissues were imaged on the SPE using a x40 (1.15NA), x63 (1.3 NA) oil objectives, and on the Nikon C2+ using a x40 (1.3 NA) oil objective.

### Quantification and statistical analysis

All image analysis was performed in FIJI/ImageJ. Time-lapse images of SOP cell signaling filopodia were acquired and lengths were measured using FiloQuant and Trackmate plugins in FIJI/ImageJ (Jacquemet et al., 2019). Number of filopodia per cell over either short windows of time or in freeze frames were quantified manually. N^sfGFP^ nuclear fluorescence was measured from 12h AP to nuclear envelope breakdown (NEBD), as previously described (Hunter et al., 2016). Briefly, an ROI was drawn in the nuclei of an N^sfGFP^ expressing cell, and the average fluorescence intensity within that ROI was reported for each time frame of the time-lapse. At the point of NEBD, a nucleus is no longer distinguishable using N^sfGFP^ and measurements are stopped. Statistical analysis was performed using GraphPad Prism. Specific statistical tests are described in the figure legends.

### Drosophila stocks

The following stocks used in this study are available at the Bloomington Drosophila Stock Center (BDSC, Bloomington, Indiana): UAS-Myosin XV RNAi, w1118, UAS-white RNAi, UAS-Myosin VIIa RNAi, UAS-Myosin VIIb RNAi. We also used the Myosin XV allele syph^21J^/FM7Ci,Act5c-GFP (Rich et al., 2021). The following GAL4 background lines were used, that were previously published elsewhere: N^sfGFP^, neuralized::H2BmRFP/CyO-GFP; neuralized-GAL4/TM6B (Hunter et al., 2016). shotgunGFP, neur-GFP-moesin^CA^/CyO-GFP; pannier-GAL4/TM6B and neuralized-GAL4, UAS-GFP-moesin^CA^/TM6B (Cohen et al., 2010). The following UAS lines were also used, that were previously constructed and published in (Rich et al., 2021): Sp/CyO; UAS-syphGFP ΔFERM(2)/TM6B, UAS-syphRFP, UAS-syphGFP ΔFERM(1)/CyOGFP, UAS-syphGFP^R213A, G434A^/CyOGFP, UAS-syphGFP/CyOGFP, UAS-syphGFP ΔMyTH4(2)/CyOGFP, UAS-syphGFP Δmotor/TM6BTb, UAS-syphGFP ΔMyTH4 (1, 2), UAS-syphGFP Δcargo, UAS-syphGFP ΔFERM(1)-ΔMyTH4(2)-ΔFERM(2), UAS-syphGFP ΔMyTH4(2)-ΔFERM(2), and UAS-syphGFP ΔMyTH4(1).

## Results

### Loss of Myosin XV leads to increased bristle density

Disruptions to the morphology or dynamics of signaling filopodia have been shown to lead to changes in the overall density and organization of bristle precursor cells at the end of bristle pattern formation (Cohen et al., 2010; Hadjivasiliou et al., 2016; Hunter et al., 2019). Specifically, conditions that generate shorter or longer signaling filopodia are predicted to lead to more or less dense bristle patterns, respectively. The unconventional MyTH4-FERM myosins function as plus-end directed motors on bundled actin structures (Sellers, 2000). This class of myosins are implicated in both trafficking of proteins along protrusions, as well as in their formation (Fitz et al., 2023; Weck et al., 2017). To determine if any of these myosins are important for signaling filopodia activity, we used an RNAi-based strategy to knockdown their expression throughout the central notum region using pannier-GAL4 (pnr-GAL4). We targeted the expression of Myosin VIIa, Myosin VIIb and Myosin XV using UAS-inducible shRNA transgenics (RNAi). The Drosophila genome does not contain Myosin X, which is the fourth major myosin of this group in mammalian organisms. The thorax bristle pattern is complete by 24 hours after pupariation (hAP). We hypothesized that pnr-GAL4 mediated knockdown of any MyTH4-FERM myosin that functions in signaling filopodia behaviors would disrupt the density of the final bristle pattern. We found that loss of Myosin XV leads to an increase in the density of bristle precursor cells by 24 hAP (Figure 1A-B).

**Fig 1.**
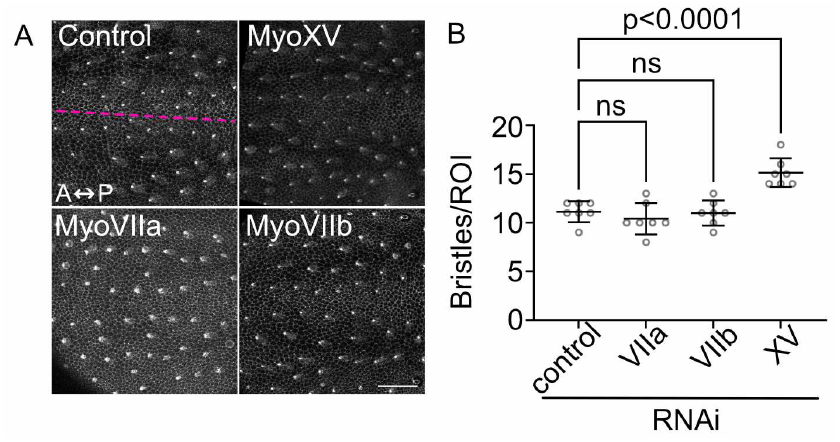
The MyTH4-FERM Myosin XV plays a role in bristle density. (A) RNAi targeting Myosin XV (MyoXV), Myosin VIIa (MyoVIIa), Myosin VIIb (MyoVIIb) or white (control) were expressed in the central notum using the pannier-GAL4 driver. Tissues also express E-cadherin GFP to label all cell boundaries and neuralized-GFP-moesin^CA^ to label bristle precursor cells. Images are from 24 hAP pupae. Scale bar, 50 μm. Anterior (A), Posterior (P). Midline is labeled in control (pink dashed line) and all images are oriented similarly. (B) Quantification of bristle density in RNAi-expressing 24 hAP pupae. One-way ANOVA with multiple comparisons (Dunnett’s) was performed. Ns = not significant. 7 ROI were analyzed across a minimum of 4 pupae for each genotype. Individual data points (circles) with mean ± SD (overlay) shown.

We did not observe any changes in the density of bristles in Myosin VIIa or VIIb RNAi expressing tissues (Figure 1A-B). Therefore, we focused on the role of Myosin XV in long-range lateral inhibition. In Drosophila, Myosin XV is an approximately 330kDa protein encoded by the gene *myo10a* which is also called *sisyphus* (*syph*). Note, 10a refers to the cytogenetic position of the gene on the X-chromosome.

### Loss of Myosin XV disrupts the development of the bristle pattern

The formation of the notum bristle pattern takes several hours over pupal stages (Cohen et al., 2010; Corson et al., 2017). We next asked how the development of pattern at earlier stages (14 hAP) in Myosin XV RNAi expressing tissues was disrupted. We visualized the selection of bristle precursor cells using neuralized promoter driven GFP-moesin^CA^, which drives an GFP-tagged moesin based F-actin reporter (GMCA; Edwards et al., 1997) in bristle precursor cells only (Figure 2A-A’). We simultaneously used pnr-GAL4 to express Myosin XV RNAi throughout the central notum epithelium. We found that decreased Myosin XV throughout the nota leads to an over-production of GFP-positive cells compared to controls (Figure 2B-B’). We also observe that the development of bristle rows 2-4 are disorganized relative to controls. Since neuralized is primarily expressed in cells that are Notch inactive, we interpret this to mean that loss of Myosin XV leads to a disruption in timely Notch signaling.

**Fig 2.**
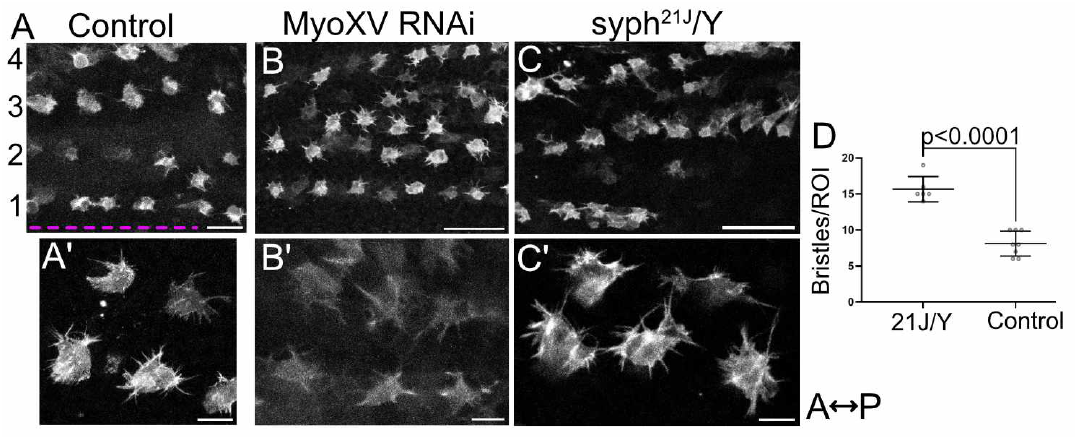
Decreased expression of Myosin XV leads to defects in bristle precursor organization during patterning. (A) Control pupae (pannier-GAL4 *>* UAS-white RNAi) expressing neuralized-GFP-moesin^CA^ to label bristle precursor cells, 14 hAP. Scale bar, 25 μm. Midline is labeled in control (pink dashed line), and all images are oriented similarly. Anterior, left; Posterior, right. Bristle rows are labelled 1-4. (A’) Control pupae with same genotype and timing as (A), zoomed in to visualize cell morphology. Scale bar 10 μm. (B) Myosin XV RNAi pupae (pannier-GAL4 *>* UAS-myosin XV RNAi) expressing neuralized-GFP-moesin^CA^ to label bristle precursor cells, 14 hAP. Scale bar, 50 μm. (B’) Pupae with same genotype and timing as (B), zoomed in to visualize cell morphology. Scale bar, 10 μm. (C) Hemizygous males carrying the syph^21J^ allele, and neuralized-GAL4 *>* UAS-GFP-moesin^CA^ to label bristle precursor cells, 14 hAP. Scale bar, 50 μm. (C’) Pupae with same genotype and timing as (C), zoomed in to visualize cell morphology. Scale bar 10 μm. (D) Quantification of 24 hAP bristle density in hemizygous male escapers carrying the syph^21J^ allele (21J/Y), compared to controls (neuralized-GAL4 *>* UAS-GFP-moesin^CA^). n = 6 ROIs across 3 21J/Y animals and 8 ROIs across 4 control animals. Individual data points (circles) with mean ± SD (overlay) shown. Significance was determined by unpaired student’s t-test.

To verify our findings with Myosin XV RNAi, we next observed patterns in pupae carrying the FRT-mediated knockout allele of Myosin XV, syph^21J^ (Rich et al., 2021). The entire coding region of Myosin XV is removed in this mutant. We co-expressed neur-GAL4, UAS-GMCA to label bristle precursor cells, imaged live male pupae hemizygous for the syph^21J^ mutation, and observed their bristle patterns at 14 hAP (Figure 2C-C’). Both RNAi knockdown and the null allele have severe developmental defects that lead to death during larval stages (data not shown)(Rich et al., 2021). Compared to control males, GFP-positive cells in syph^21J^mutant pupae are often grouped along rows rather than isolated (Figure 2C). Those cells that are isolated are closer to each other than controls (21.0 ± 8.0 μm across n =55 syph^21J^ cell pairs, compared to 29.3 ± 8.4 μm across n = 43 control cell pairs; p*<*0.0001 by student’s t-test). Consistent with the observation that knockdown of Myosin XV by RNAi leads to increased bristle density, hemizygous syph^21J^ pupae also have increased bristle density relative to controls (Figure 2D). The signaling filopodia of syph^21J^ mutant cells appear indistinguishable from wildtype (Figure 2C’). In contrast to Myosin XV RNAi expressing pupae, we find that syph^21J^ pupae sometimes have developmental delays such that entire bristle rows have not yet appeared by 14 hAP (Figure 2C). Altogether, our analysis of tissue-wide RNAi-mediated knockdown or a null allele of Myosin XV show that wildtype Myosin XV plays a role in the long-range lateral inhibition mechanism that regulates the well-spaced array of sensory bristles on the dorsal thorax.

### Decreased expression of Myosin XV in bristle precursors increases filopodia length and number

The spacing of bristle precursor cells depends on a dynamic length determinant, in part facilitated by the dynamics of basal signaling filopodia (Cohen et al., 2010; Corson et al., 2017; Hadjivasiliou et al., 2016). We next asked whether loss of Myosin XV caused defects specifically associated with the morphology or behavior of signaling filopodia. Based on our quantification of bristle density, we hypothesized that decreased Myosin XV expression leads to shorter and fewer signaling filopodia. Since syph^21J^ mutants and pnr-GAL4 driven RNAi pupae do not produce large amounts of samples due to increased lethality at earlier stages, we next used neuralized-GAL4 to express UAS-Myosin XV RNAi in bristle precursor cells only, while co-expressing UAS-GMCA to visualize cell shape and filamentous actin structures. We observe that signaling filopodia in bristle precursor cells expressing Myosin XV RNAi appear morphologically indistinguishable from control signaling filopodia (Figure 3A-B).

**Fig 3.**
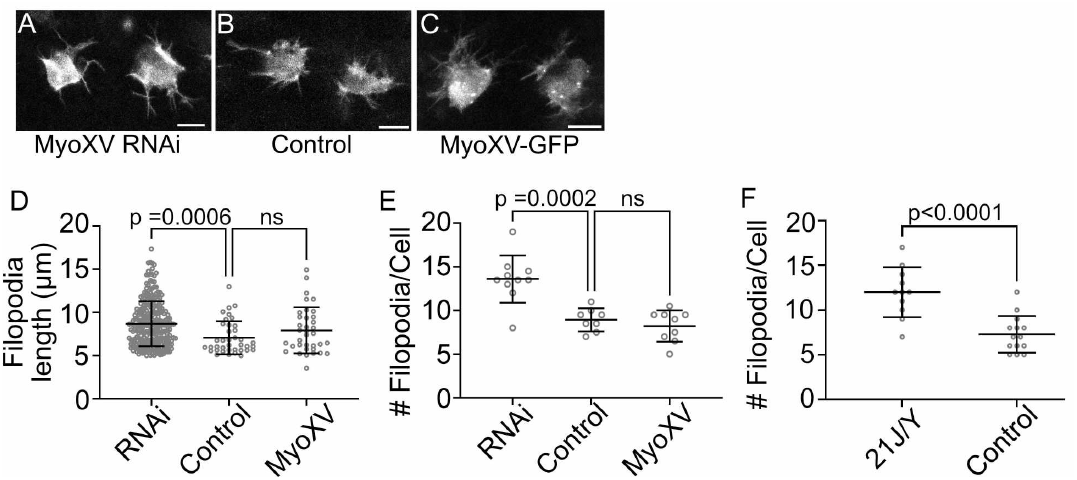
Myosin XV negatively regulates signaling filopodia length and number. (A-C) Stills from movies used to quantify data in (D-E). (A) Myosin XV RNAi = neur-GAL4 *>* UAS-GMCA, UAS-Myosin XV RNAi pupae, at 14 hAP. (B) Control = neur-GAL4 *>* UAS-GMCA, UAS-white RNAi pupae, at 14 hAP. (C) Myosin XV GFP = neur-GAL4 *>* UAS-GMCA, UAS-GFP full length Myosin XV pupae, at 14h AP. Scale bars on all images, 10 μm. (D) Quantification of filopodia length according to genotypes as in (A-C). (E) Quantification of filopodia produced per cell over 15 minutes, according to genotypes as in (A-C). One-way ANOVA with multiple comparisons (Dunnett’s) was performed. Ns = not significant. (F) Quantification of filopodia per cell (1 frame), in hemizygous syph^21J^ and control pupae. n = 11 cells across 3 21J/Y animals and 14 cells across 3 control animals. Significance was determined by unpaired student’s t-test. For (D-F), individual data points (circles) with mean ± SD (overlay) shown. Minimum of 3 pupae were imaged and analyzed per genotype.

To quantify the formation and dynamics of signaling filopodia over time, we used FiloQuant to automatically track and measure filopodia in bristle precursor cells (Jacquemet et al., 2019). We found that signaling filopodia in Myosin XV knockdown cells are longer (8.7 ± 2.6 μm, n = 272 filopodia) compared to control cells (7.1 ± 1.9 μm, n = 38 filopodia)(Figure 3D). We also observed more filopodia in Myosin XV knockdown cells (27.2 ± 5.4 filopodia per cell, n = 10 cells, N = 4 pupae) compared to control cells (17.9 ± 2.6 filopodia per cell, n = 8 cells, N = 3 pupae) (Figure 3E). Consistent with this result, at 14 hAP syph^21J^ mutant bristle precursors also have more filopodia (12.0 ± 2.8 filopodia per cell, n = 11 cells, N = 3 pupae) relative to controls (7.3 ± 2.1 filopodia per cell, n = 14 cells, N = 3 pupae)(Figure 3F) Given these results, we hypothesized that over-expression of Myosin XV could be sufficient to generate fewer or shorter filopodia. Surprisingly, we found that over-expression of full length Myosin XV does not change the length (7.9 ± 2.7 μm, n = 36 filopodia) or number of signaling filopodia (16.4 ± 3.6 filopodia per cell, n = 9 cells, N = 3 pupae) (Figure 3C-E). Taken together, these results suggest that Myosin XV is a negative regulator of signaling filopodia length and formation.

### Myosin XV localizes along the lengths of signaling filopodia

Our results thus far are consistent with Myosin XV playing a role in the formation and dynamics of signaling filopodia during notum patterning. We next wanted to determine the sub-cellular localization of Myosin XV in bristle precursor cells. Mammalian Myosin XV plays a major role in the formation and maintenance of mechanosensory stereocilia and localizes to the tips of these bundled actin structures (Belyantseva et al., 2005; Manor et al., 2011). In elongating sensory bristles (≈33 hAP), Drosophila Myosin XV localizes along the length and at the tips of developing bristles (Rich et al., 2021). In embryos and in insect cell culture, Myosin XV localizes along the length and at the tips of filopodia (Liu et al., 2007). Since no antibody currently exists to visualize endogenous Drosophila Myosin XV, we instead over-expressed RFP-tagged full length Myosin XV in live bristle precursor cells. We co-expressed GMCA in these cells to simultaneously visualize F-actin. We observed localization of Myosin XV along signaling filopodia, as well as their tips (arrowheads, Figure 4). This result is consistent with our knockdown data, suggesting that Myosin XV plays a role in signaling filopodia dynamics.

**Fig 4.**
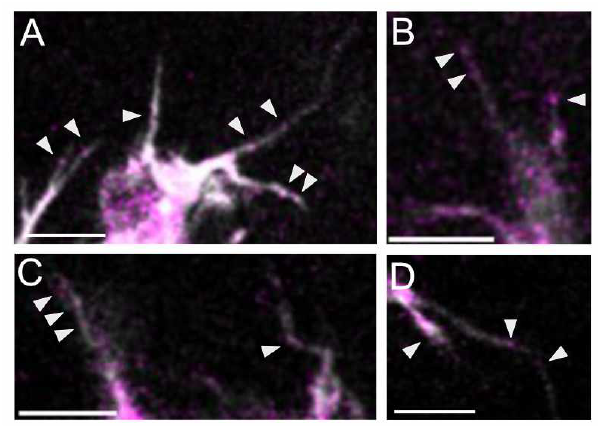
Localization of Myosin XV in bristle precursor cell signaling filopodia. (A-D) Stills from four individual bristle precursor cells across 3 pupae expressing neuralized-GAL4 *>* UAS-GMCA, UAS-RFP-Myosin XV. Scale bars on all images, 5 μm. Arrowheads indicate RFP puncta within a cellular projection.

### Decreased expression of Myosin XV dampens Notch signaling activity

The specification of bristle precursor cells in the initially unpatterned notum is driven by Notch-mediated lateral inhibition. Long-range Notch signaling is facilitated by the activity of signaling filopodia. Based on our results showing that decreased Myosin XV leads to altered filopodia length and number, and that the overall patterning is altered in these tissues, we hypothesized that decreased levels of Myosin XV in signal sending cells (bristle precursors) would disrupt Notch activation in neighboring epithelial cells, relative to controls. To test this hypothesis, we expressed Myosin XV RNAi in bristle cells only using neuralized-GAL4, in a notum epithelium that expresses the N^sfGFP^ Notch transcriptional reporter (Figure 5A). This reporter expresses a nuclear localized, PEST-tagged, superfolder GFP under the control of a Notch responsive promoter (Hunter et al., 2016). We then tracked sfGFP fluorescence in nuclei of cells adjacent to or one cell diameter distant from the nearest bristle precursor cell. We term these cells adjacent or distant epithelial cells, respectively (see Figure 5B-B’ insets). Adjacent cells can engage in Notch signaling with a bristle precursor via both large cell-cell interfaces and signaling filopodia. In contrast, distant cells can only engage in Notch signaling with a bristle precursor via signaling filopodia. Importantly, by using neuralized-GAL4, all adjacent and distant epithelial cells express wildtype levels of Myosin XV; only bristle precursor cells see Myosin XV knockdown. We tracked nuclear fluorescence levels from 12 hAP until nuclear envelope breakdown (NEBD), which occurs between 15-20 hAP. Loss of Myosin XV in bristle cell precursors alone leads to a decreased Notch response in all wildtype neighboring epithelial cells (Figure 5B-B’). These results are consistent with our observation of increased GFP-positive cells during patterning stages (Figure 2B), as an increase in cells with activation of neuralized expression indicates a decrease in Notch activation. Together, this data shows that Myosin XV plays role in the activation of robust Notch signaling both in adjacent and distant cells, both of which may signal to bristle precursor cells via signaling filopodia.

**Fig 5.**
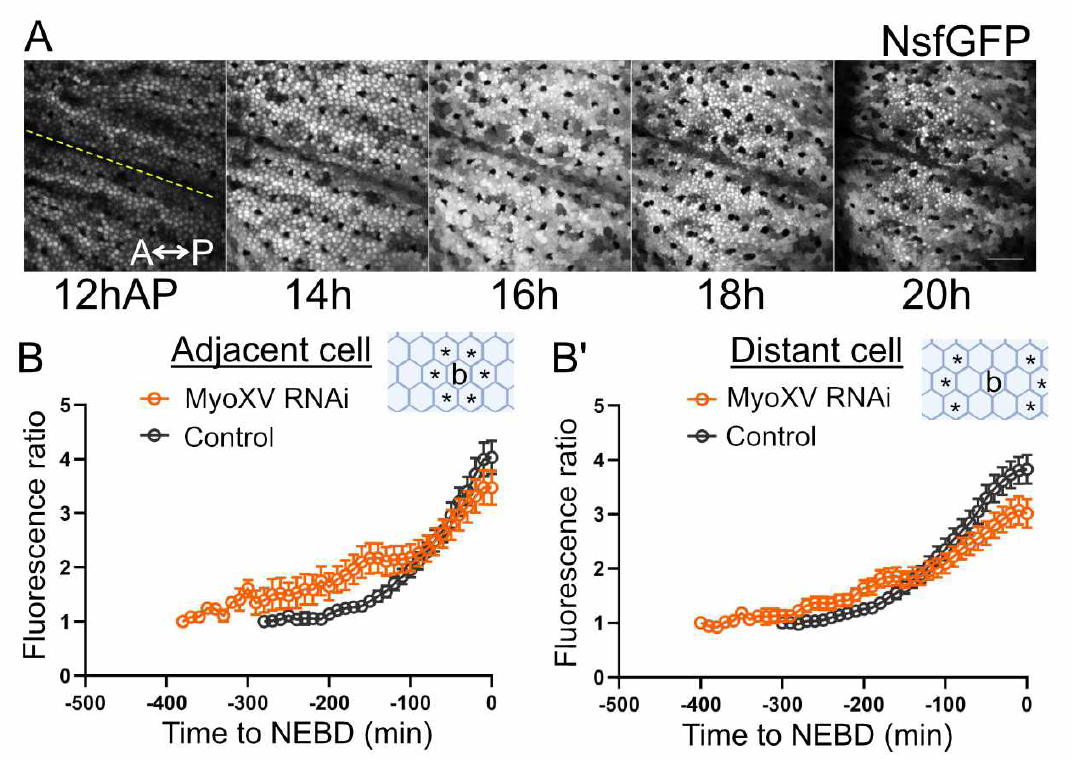
Myosin XV is required for robust Notch signaling response. (A) Montage from a control pupae expressing the Notch transcriptional reporter N^sfGFP^(grayscale). Anterior, left; Posterior, right. Yellow line in 12h AP image indicates the animal midline. Notch response in epithelial cells peaks 14-16 hAP, immediately followed by mitosis 16-18 hAP, and termination of Notch signaling by 20-24 hAP. Scale bar, 50 μm. (B-B’) Nuclear GFP fluorescence in signal receiving cells (B) adjacent to bristle precursor cells or (B’) distant from bristle precursor cells. Inset cartoons illustrate cell position (*) relative to bristle precursor cell (b) for each group. GFP signal is normalized to initial fluorescence level. All time-lapse measurements were aligned such that time = 0 minutes at nuclear envelope breakdown (entry into mitosis). Myosin XV RNAi (orange): 50 nuclei of each position (adjacent or distant) were measured across 3 pupae. Control (black): 20 nuclei of each position (adjacent or distant) were measured across 3 pupae. Mean ± SEM shown.

### Myosin XV plays a role in the localization of Delta and Notch to signaling filopodia

Our results support a model in which Myosin XV indirectly supports long-range signaling through the regulation of filopodia number and length. However, it remains unclear why the formation of more and longer signaling filopodia would lead to a decrease in Notch response in signal receiving cells, and increased bristle density. We hypothesized that although more signaling filopodia are being formed in Myosin XV RNAi expressing cells, these signaling filopodia are not necessarily competent to signal. In other filopodia and stereocilia, Myosin XV plays a role in trafficking proteins along bundled actin structures towards the tip (Belyantseva et al., 2005; Manor et al., 2011; Weck et al., 2017). It is currently unknown if Myosin XV directly interacts with Notch or Delta.

To address our hypothesis, we investigated the levels and localization of Delta ligand and Notch receptor in cells with wildtype or decreased expression of Myosin XV. In the notum, apical Delta and Notch are localized primarily to the apical cell-cell junctions (Bellec et al., 2020; Benhra et al., 2010). Cytoplasmic puncta staining positive for Delta or Notch can also be observed in bristle precursor cells. The cytoplasmic puncta may represent: (1) endocytosed Delta ligand and trans-endocytosed Notch extracellular domain (NECD) post-cleavage, or (2) Delta and Notch being trafficked to the bristle precursor cell surface. Additionally, all cells in the notum are expected to express at least low levels of both ligand and receptor (Collier et al., 1996). On the basal surface, Delta and Notch puncta are associated with signaling filopodia, as discrete puncta, often at the tips of signaling filopodia (Figure 6A-B) (Hunter et al., 2019). In control cells, we observe Delta positive puncta within the cell body and, to a lesser extent, within filopodia (Figure 6A). NECD positive puncta are also visible both in the cell body and within filopodia (Figure 6B). We do not observe any difference between the number of Delta (5.2 ± 2.0 puncta, n = 11 cells) and Notch (5.1 ± 2.3 puncta, n = 34 cells; n.s. by student’s t-test) positive puncta in control bristle precursor cells. Importantly, this result does not distinguish whether the Delta or Notch positive puncta are a result of signaling between adjacent neighbors or distant neighbors.

**Fig 6.**
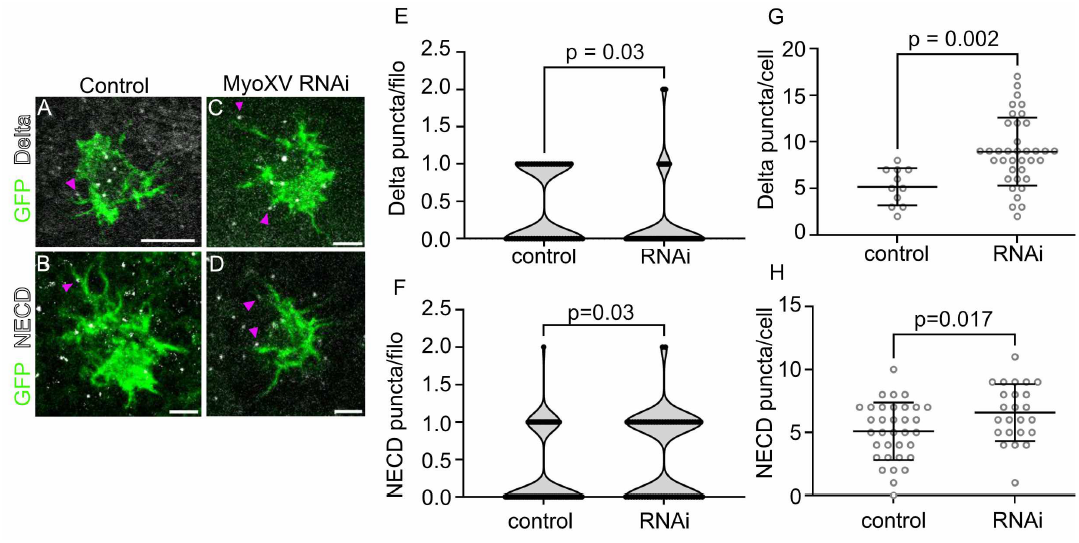
Myosin XV is required for the balance of Delta and Notch in filopodia. (A-D) Example of fixed images used to generate data in (E-H). Control pupae of the genotype neur-GAL4 *>* UAS-GMCA, UAS white RNAi stained with anti-GFP (green) and (A) anti-Delta antibody (white) or (B) anti-Notch extracellular domain (NECD, white). Myosin XV knockdown pupae of the genotype neur-GAL4 *>* UAS-GMCA, UAS Myosin XV RNAi stained with anti-GFP (green) and (C) anti-Delta antibody (white) or (D) anti-NECD (white). Magenta arrowheads indicate Delta or NECD puncta on signaling filopodia. Scale bars for all images, 5 μm. (E) Quantification of Delta puncta on signaling filopodia in control (0.42 ± 0.5 puncta, n = 48 filopodia in 13 cells across 4 pupae) and Myosin XV RNAi (0.21 ± 0.5 puncta, n = 72 filopodia in 16 cells across 4 pupae) expressing bristle precursor cells. (F) Quantification of NECD puncta on signaling filopodia in control (0.32 ± 0.5 puncta, n = 62 filopodia in 15 cells across 4 pupae) and Myosin XV RNAi (0.54 ± 0.6 puncta, n = 59 filopodia in 17 cells across 4 pupae) expressing bristle precursor cells. (G) Total Delta puncta per cell in control (5.2 ± 2.0 puncta, n = 11 cells across 4 pupae) and Myosin XV RNAi (8.9 ± 3.6 puncta, n = 37 cells across 11 pupae) expressing bristle precursor cells. (H) Total NECD puncta per cell in control (5.1 ± 2.3 puncta, n = 34 cells across 12 pupae) and Myosin XV RNAi (6.6 ± 2.3 puncta, n = 24 cells across 8 pupae) expressing bristle precursor cells. For (E-H), individual data points shown (circles). Mean ± SD is overlain in (G-H). Mean ± SD reported in legend above, and significance was determined by unpaired student’s t-test.

To determine if the expression level of Myosin XV plays a role in the localization of ligand and receptor in bristle precursor cells, we performed immunofluorescence staining for Delta or NECD in tissues with bristle precursor cells labeled with GFP (neuralized-GAL4, UAS-GMCA) and co-expressing UAS-Myosin XV RNAi (Figure 6C-D). Consistent with our live filopodia tracking results, we observe an increase in the average number of signaling filopodia in bristle precursor cells expressing Myosin XV RNAi (8.6 ± 2.7 filopodia per cell, n = 61 cells) compared to controls (6.5 ± 2.1 filopodia per cell, n = 45 cells; p *<* 0.0001 by student’s t-test). Signaling filopodia were also longer in Myosin XV RNAi cells (7.0 ± 2.6 μm, n = 110 filopodia) compared to controls (6.4 ± 1.9 μm, n = 131 filopodia; p = 0.03 by student’s t-test), although both were shorter than measured in live cells, which may be in part due to fixation methods.

Bristle precursor cells expressing Myosin XV RNAi exhibited a misbalance in Delta and NECD positive puncta throughout the cell body and filopodia (Figure 6E-H). First, we analyzed the number of Delta or Notch puncta along signaling filopodia. We observe fewer Delta positive puncta in Myosin XV RNAi filopodia compared to controls (Figure 6E), and more Notch extracellular domain positive puncta in Myosin XV RNAi filopodia compared to controls (Figure 6F). We find that control filopodia contain roughly the equal numbers of Delta and NECD puncta along their lengths (n.s. by student’s t-test). However we find that Delta puncta are underrepresented in Myosin XV RNAi filopodia compared to NECD puncta in the same cell genotype (p=0.0005 by student’s t-test). Next we compared the levels of Delta and NECD puncta throughout the cell body (including filopodia). Myosin XV RNAi expressing cells exhibit more Delta or NECD positive puncta throughout the cell body compared to controls (Figure 6G-H). Control bristle cells show no significant difference (student’s t-test) between number of Delta or NECD puncta throughout the cells, that is, control cells have equal amounts of Delta and NECD puncta throughout the cell body. We observe that Delta puncta are over-represented in the Myosin XV RNAi expressing cell body compared to NECD positive puncta (p = 0.006, student’s t-test). These data suggest that the balance of Delta ligand mediated Notch activation is disrupted in bristle precursor cells that express Myosin XV RNAi. The finding that there is decreased Delta puncta on filopodia is also consistent with our findings that Notch signaling is decreased in neighboring epithelial cells. Our interpretation of these data are that decreased Myosin XV expression leads to the formation of increased, but not necessarily signaling competent, signaling filopodia.

### Myosin XV motor activity is required to promote signaling filopodia formation

Finally, we investigated how Myosin XV regulates both the morphology of signaling filopodia and the localization of Delta. Full length Myosin XV comprises the following domains: motor, IQ motifs, two FERM (band 4.1, ezrin, radixin, moesin) domains and two MyTH4 (Myosin tail homology 4) domains (Rich et al., 2021)(Figure 7A). Notably, Drosophila Myosin XV lacks the large N-terminal extension found in mammalian Myo15. In order to determine the domains of Myosin XV are required for signaling filopodia formation, we over-expressed GFP-tagged truncated Myosin XV constructs in bristle precursor cells expressing neur-GAL4, UAS-GMCA. We analyzed the number of filopodia formed by a bristle precursor cell in pupae at 14 hAP. Over-expression of the domain deletion constructs does not lead to any gross morphological defects in bristle precursor cell shape or the ability to form basal cytonemes (Figure S1). We did however observe that expression of the Δmotor construct leads to nuclear GFP localization (Figure 7C). Despite this, Δmotor expressing cells are still able to form signaling filopodia, perhaps due to the presence of endogenously expressed Myosin XV. When we analyzed the number of filopodia formed by cells over-expressing different Myosin XV deletion domain constructs, we found that only expression of the point mutant Myosin XV^R213A, G434A^ led to the overproduction of signaling filopodia compared to control cells over-expressing full length Myosin XV (Figure 7D-E). Myosin XV^R213A, G434A^ is a motor dead construct that changes key amino acid residues in switch I and II of the head domain to Alanine (Rich et al., 2021). This construct may bind to filamentous actin, but is unable to move along filaments. We did not observe any nuclear GFP signal in Myosin XV^R213A, G434A^ expressing cells, which may partially explain why the two conditions have different phenotypes. Therefore, this finding is consistent with our data showing that loss of Myosin XV expression increases filopodia number (Figure 3E-F). Over-expression of Myosin XV truncation mutants lacking all or parts of the cargo domain did not affect the number of filopodia formed (Figure 7E).

**Fig 7.**
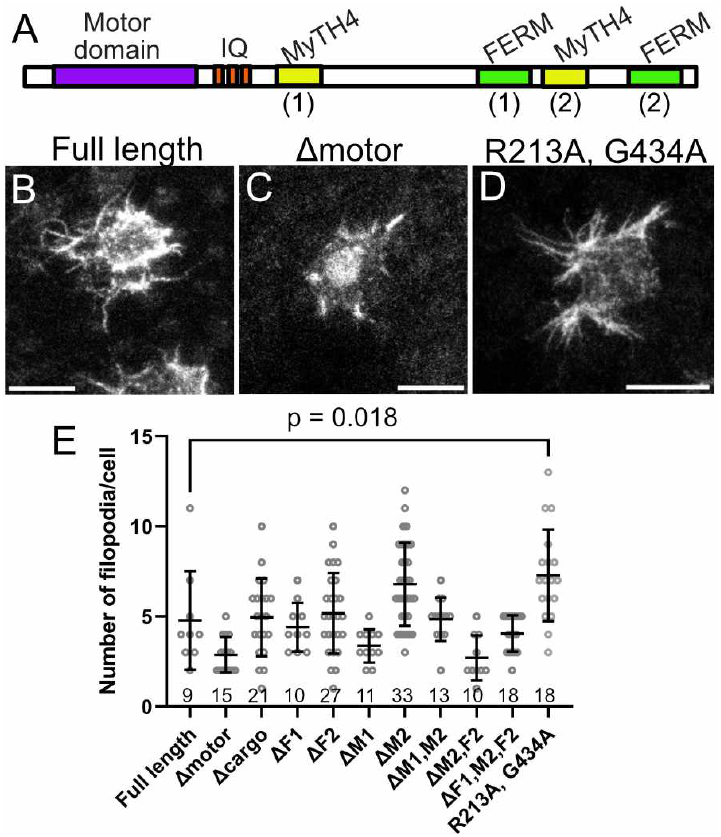
Myosin XV motor activity is required for its effects on filopodia length. (A) Organization of Myosin XV protein domains. (B-D) Example images used to generate data in (D). See Supplement Figure S1 for additional panels. Neuralized-GAL4 was used to co-express GFP tagged deletion domain or mutant constructs listed, e.g., neur-GAL4 *>* UAS-GMCA, UAS-GFP-Δmotor-Myosin XV. Pupae are stained with anti-GFP antibody. Scale bars for all images, 5 μm. (E) Number of filopodia per bristle precursor cell overexpressing the listed Myosin XV construct was quantified in fixed tissues. n = number of cells analyzed, in a minimum of 3 pupae per genotype. Individual data points shown (circles), with mean ± SD overlay. One-way ANOVA with multiple comparisons (Dunnett’s) was performed. Individual comparisons to full length Myosin XV were n.s. unless indicated.

We also analyzed whether over-expression of domain deletion constructs could disrupt the number of Delta puncta in bristle precursor cell filopodia. We hypothesized that loss of specific cargo domains would lead to decreased Delta puncta in filopodia due to failure to form trafficking complexes. However, we did not observe any differences in the number of Delta puncta along filopodia relative to filopodia in cells expressing full length Myosin XV control (Figure S1). One possibility for this result is the presence of sufficient endogenous Myosin XV to traffick cargo to or within filopodia. Altogether our results with the domain deletion constructs indicate that the motor activity of Myosin XV is required for the negative regulation of signaling filopodia.

## Discussion

Cell morphology plays an important role in the ability of cells in tissues to send and receive signals during development. The formation and activity of cytonemes, or signaling filopodia, are particularly interesting as they play an important role in the active distribution of morphogens, and have been implicated in the development of several tissues in vertebrates and invertebrates (Kornberg, 2017). Therefore it is important to understand how changes in cell morphology affect the ability of cells to send or receive signal. Here we identified a role for the Myosin XV in the regulation of signaling filopodia dynamics and Notch mediated lateral inhibition during bristle patterning.

Of the three unconventional myosins present in the Drosophila genome, we identified Myosin XV as a key regulator of bristle pattern density. Loss of Myosin XV expression led to an increased bristle density, whereas loss of Myosin VIIa and VIIb did not. Based on previous mathematical models (Cohen et al., 2010; Hadjivasiliou et al., 2016), we anticipated that loss of Myosin XV was associated with shorter signaling filopodia because decreasing the length scale of Notch signaling leads to patterns that are more dense. We were then surprised to find that Myosin XV negatively regulates filopodia length and number in bristle precursor cells during lateral inhibition stages. How might decreased levels of Myosin XV lead to longer and increased numbers of signaling filopodia? The best studied Myosin XV is mammalian Myo15, which localizes to stereocilia in hair cells of the cochlea (Belyantseva et al., 2003; Fang et al., 2015).

Myo15, along with its binding partners Whirlin and Eps8, is associated with the plus-ends of the bundled actin filaments at the tips of stereocilia (Belyantseva et al., 2005; Manor et al., 2011; Mauriac et al., 2017; Moreland et al., 2021). Loss of Myo15 is associated with shorter and increased numbers of stereocilia structures (Moreland et al., 2021). Why loss of Myo15 leads to increased numbers of stereocilia is unclear. The effect of Myosin XV on the elongation of signaling filopodia could be direct or indirect. Mammalian Myo15 has been shown to directly participate in the nucleation of F-actin filaments in vitro (Gong et al., 2022) and the association of the Myo15 motor domain with the plasma membrane is sufficient to stimulate the formation of filopodia in cell culture (Fitz et al., 2023). It is currently unknown if Drosophila Myosin XV has a similar ability. In that case however, we would expect that loss of Myosin XV would shorten signaling filopodia. A recent study showed that Drosophila Myosin XV associates with MICAL, an enzyme that promotes the disassembly of filamentous actin in elongating bristles (Rich et al., 2021). If Myosin XV were involved in the trafficking of negative regulators to the tips of the filamentous actin bundles inside cytonemes, then we might expect to see increased length when Myosin XV levels are decreased. Currently, we do not know what proteins interact with Myosin XV in the patterning notum.

If Myosin XV is a negative regulator of signaling filopodia and length, then why does decreased Myosin XV expression result in a bristle pattern with increased bristle density compared to controls? Our data indicate that that although more and longer signaling filopodia are present, they may not be competent to signal. Several studies support a role for signaling filopodia in long-range Notch signaling (Cohen et al., 2010; de Joussineau et al., 2003; Hadjivasiliou et al., 2016). Non-adjacent cells contact each other using filopodia, which have been shown to carry both Notch receptor and Delta ligand (Hunter et al., 2019; Renaud and Simpson, 2001). Drosophila Myosin XV is not known to directly interact with either Delta or Notch. However, a yeast two-hybrid assay for potential interactors of the Myosin XV C-terminal tail identified Nedd4, an E3 ubiquitin ligase that is required for the internalization and inactivation of Notch (Liu et al., 2007; Sakata et al., 2004). Our data shows that there are fewer Delta puncta but more NECD puncta per filopodia in bristle precursor cells. Since bristle precursor cells are Notch inactive, but are thought to have expressed low levels of Notch (Collier et al., 1996), one possibility is that decreased Myosin XV levels leads to failure to down-regulate Notch in bristle precursor cell signaling filopodia. Cis-interactions with increased levels of Notch along filopodia and remaining Delta along filopodia could lead to lower levels of Notch signaling in receiving cells due to decreased available surface ligand (Sprinzak et al., 2011, 2010). Lowering the level of Notch activation in distant cells, in turn, increases the possibility that those cells will undergo a cell fate switching event. This would lead to the selection of too many bristle precursor cells, which is consistent with our findings that (1) lowered Myosin XV expression leads to a decreased Notch response in epithelial cells distant from the bristle precursor cell, and (2) bristle precursor cells are overproduced during active patterning stages.

In bristle precursor cells, we found that Myosin XV localizes at the tips and along the length of signaling filopodia. The C-terminal tail of MyTH4-FERM domain containing myosins is known to be important for interactions with cargo, cytoskeleton, and membrane (Weck et al., 2017). The interactions in the tail domains impact the ability of a myosin to interact with, and process along, bundled filamentous actin. We were interested in determining what domains of Myosin XV were essential for the formation of signaling filopodia and protein localization along their length. When we over-expressed Myosin XV truncation constructs in bristle precursor cells, we did not observe changes to levels of Delta puncta along filopodia relative to full length Myosin One possibility is that endogenous Myosin XV is able to traffick despite over-expression of truncation constructs. It is not currently known if Myosin XV can, or needs to, dimerize, like Myosin X (Lu et al., 2012) and other myosins. If Myosin XV dimerizes, we might expect some of the truncation constructs, especially Δmotor, to act as dominant negatives, similar to motor-less non-muscle myosin II (Franke et al., 2005). Interestingly, previous work with these truncation constructs suggested that over-expression of cargo-binding mutant forms of Myosin XV either have an over-expression phenotype or function as dominant negatives during the extension of sensory bristles (Rich et al., 2021). In future work, truncation constructs could be expressed in Myosin XV mutant tissue clones, given the larval stage lethality associated with syph^21J^ and RNAi-knockdown. A second possibility is that a separate system is responsible for distributing Delta into signaling filopodia. Microtubules have been observed to extend into filopodia in cell culture and during axon guidance (Dent et al., 2007; Schober et al., 2007), where they can influence filopodia movement. The presence of microtubules and tip directed kinesin motors within signaling filopodia would be an alternative to trafficking along bundled actin by tip directed Myosin motors. Previous findings indicate that microtubules are dispensable for the formation of signaling filopodia by bristle precursor cells, since treatment with the microtubule inhibitor colchicine does not disrupt protrusion formation (Georgiou and Baum, 2010). However this does not rule out a role for microtubules in the distribution of morphogens for signaling. Further investigations will be needed to determine how Notch and Delta proteins are targeted to, and trafficked along, signaling filopodia for long-range lateral inhibition.

Signaling filopodia are an essential active cell mechanism of cell-cell signaling, and have been shown to be important for several signaling paradigms, across many different developing tissues, in both vertebrates and invertebrates. Despite the numerous examples of the role of cytonemes in development, we still do not fully understand how these structures are formed and regulated. Here, we have shown that Myosin XV plays a role in the formation of signaling filopodia during lateral inhibition in patterning epithelia, and that it is required for organization of Delta and Notch within the cell. The mechanisms by which Myosin XV regulates signaling filopodia dynamics will be the focus of future research. There is still much to learn about the organization and dynamics of bundled actin filaments within cytonemes, the interactions of the projections with their environment, and how this all contributes to the properties that allow cytonemes to achieve their lengths and signaling specificity.

## Supporting information

**S1 Fig Overexpression of Myosin XV deletion constructs in bristle precursor cells (related to Figure 7)**.

## Author Contributions

The project was conceptualized by GH. Experiments and analysis were carried out by RC, TS, LC, JZ, AS, AC, and GH. Figures were generated by TS and GH. Original draft was written by GH. Reviewing and editing was carried out by all authors.

## Acknowledgments

We thank the Terman Lab at UTSW for sharing Drosophila reagents. We thank Ed Giniger (NINDS/NIH) for postdoctoral support. LC and JZ were funded by the McNair Scholars program. Stocks obtained from the Bloomington Drosophila Stock Center (NIH P40OD018537) were used in this study. Monoclonal antibodies (as described in Methods) were obtained from the Developmental Studies Hybridoma Bank, created by the NICHD of the NIH and maintained at The University of Iowa, Department of Biology, Iowa City, IA 52242. Work in the Hunter lab is supported by Clarkson University and the National Institutes of Health (R03NS130395) to GH. We thank Hunter lab members for their critical reading of this manuscript and comments.

