## Supplemental Figure 1 for "Myosin XV is a negative regulator of signaling filopodia during long-range lateral inhibition"

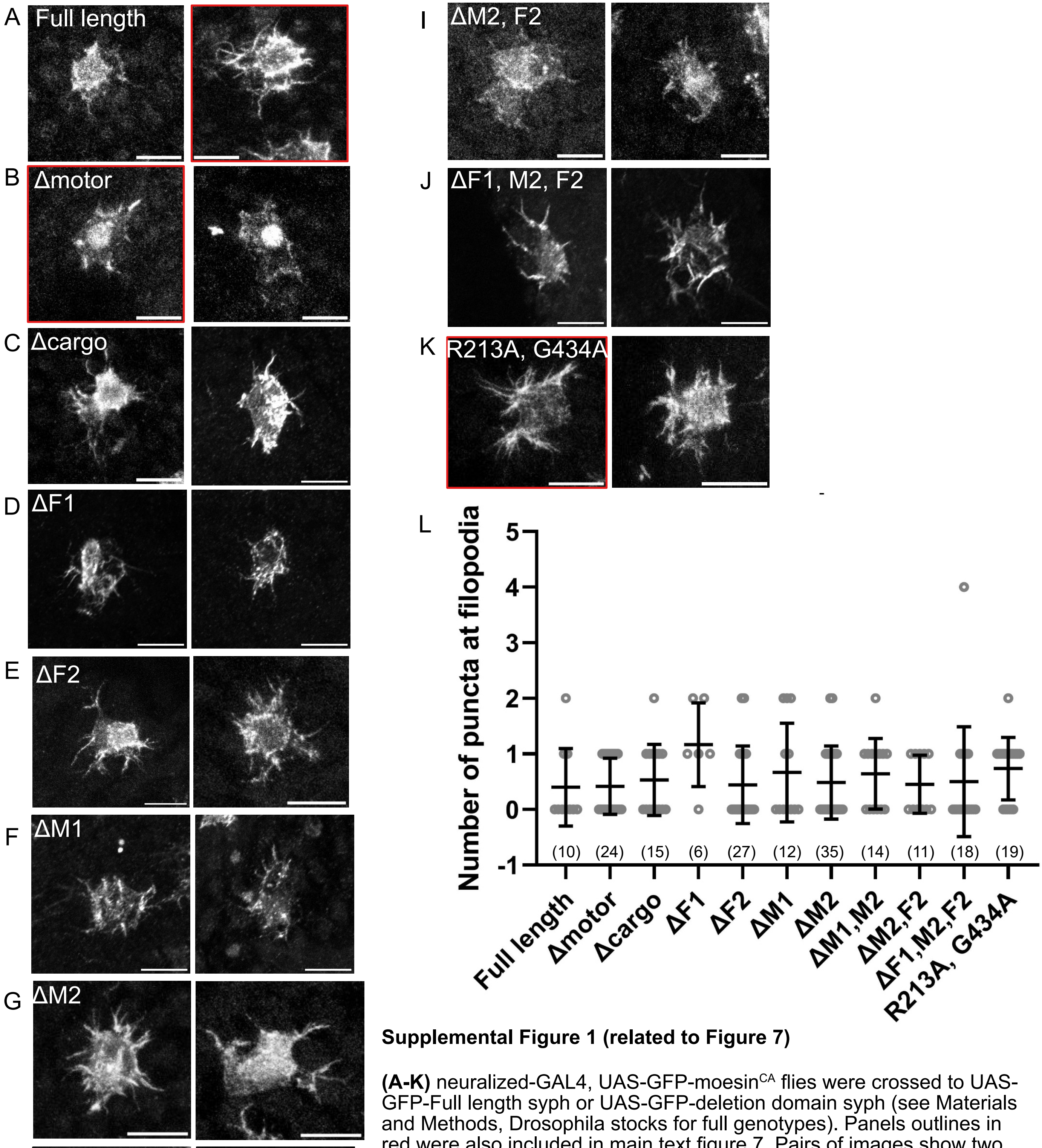


**Supplemental Figure 1 (related to Figure 7)**

**(A-K)** neuralized-GAL4, UAS-GFP-moesin$^{CA}$ flies were crossed to UAS-GFP-Full length syph or UAS-GFP-deletion domain syph (see Materials and Methods, Drosophila stocks for full genotypes). Panels outlines in red were also included in main text figure 7. Pairs of images show two example bristle precursor cells. Scale bars, 5 microns. **(L)** Quantification of Delta puncta along signaling filopodia in cells of the indicated genotype. Individual data points shown (circles), with mean ± SD overlay. One-way ANOVA with multiple comparisons (Dunnett's) was performed. Individual comparisons to full length Myosin XV were n.s. For each genotype, (n) = number of filopodia analyzed, across a minimum of 3 pupae.
